# Distinct Spatial Immune Architectures in Tumor and Tumor-Adjacent Tissues of Early-Stage Non-Small Cell Lung Cancer

**DOI:** 10.64898/2026.09.08.750122

**Authors:** Isabella Polic, Claudio Arrechedera, Jared Slone, Amanda Montoya, Emily Bontekoe, Peixin Jiang, Anika Patel, Tatiana Karpinets, Xiaogang Wu, Xingzhi Song, Qianyun Luo, Heladio Ibarguen, Auriole Tamegnon, Mei Jiang, Cara Haymaker, Annikka Weissferdt, Ara A. Vaporciyan, Lydia Kavraki, Tina Cascone, Ignacio Wistuba, ICON Study Team, Luisa M. Solis Soto, John V. Heymach, Jianjun Zhang, Don L. Gibbons, Alexandre Reuben

## Abstract

**Background:** Lung cancer remains the leading cause of cancer-related deaths in the United States with over 124,000 estimated deaths for 2026. Previous studies have found that tumor-adjacent lung tissues may provide additional insight into the immune microenvironment of early-stage NSCLC.

**Methods:** Multiplex immunofluorescence (mIF) imaging was performed on 192 tissues from 101 early-stage non-small cell lung cancer patients including 91 matched tumor-adjacent pairs, using three mIF panels. Spatial analyses were performed to identify distinctions between tissues and identify associations with clinical and genomic features.

**Results:** Tumor tissue showed significantly higher densities of T cells, macrophages, B cells, and memory/regulatory populations than adjacent tissue (p<0.001). Tumors exhibited greater spatial heterogeneity, with higher prevalence of spatial patterning (47.7% vs. 31.2%) and more consistently localized organization, whereas adjacent tissue showed stronger individual-cell clustering. Pairwise colocalization (Ripley’s L-cross) revealed selective spatial segregation in tumors (including B-cells from memory/regulatory cells and among cytotoxic T-cell subsets) and reduced immune proximity to malignant cells relative to the strong immune-epithelial association in adjacent tissue. Spatial features were linked to genomic features or patient outcomes: tumor neoantigen burden exhibited a positive association with CD3+ T cells in the tumor, while colocalization between CD45RO^+^CD57^+^GZMB^+^ cells and CD57^+^GZMB^+^ cells was negatively associated with overall survival outside the tumor.

**Conclusion:** Tumor and adjacent tissues harbor distinct spatial immune architectures, with spatial features displaying associations with genomic features or patient outcomes. These findings highlight immune microenvironment reorganization and the importance of incorporating spatial context from both compartments into risk assessment in NSCLC.

## 1 Introduction

Lung cancer is the third most commonly diagnosed cancer in the United States and remains the leading cause of cancer-related death, with 124,990 deaths projected in 2026 (1,2). Although incidence and mortality rates have declined in recent years, 29% of cancer cases are identified at a localized stage (1,2). Non-small cell lung cancer (NSCLC) accounts for approximately 85% of lung cancer cases, which includes the histological subtypes adenocarcinoma, squamous cell carcinoma, and large cell carcinoma (3). Treatment strategies vary by cancer stage and molecular characteristics, with early-stage cancer typically managed by surgical resection and advanced-stage cancer treated with targeted therapies and immunotherapies (4,5). The effectiveness of these treatments can be strongly influenced by the tumor microenvironment (TME), a complex ecosystem of cancer cells, immune cells, stromal cells, and signaling molecules that interact to regulate tumor growth, immune responses, and therapeutic outcomes (6,7). As a result, understanding the composition and organization of the TME has become increasingly important for identifying biomarkers of prognosis and treatment response.

Spatial biology has emerged as a powerful approach for investigating the TME because it preserves tissue architecture while enabling characterization of the location and interactions of cells within their native environment (8). Unlike conventional analyses that quantify cell populations without regard to their physical relationships, spatial approaches can reveal how the organization of cells and interactions within tissues influences tumor progression and immune function. In NSCLC, spatial analyses have been used to characterize tumor heterogeneity, elucidate mechanisms of immunotherapy response, and improve understanding of TME dynamics (9–12).

While most studies have focused on the tumor itself, increasing evidence suggests that tumor-adjacent tissues also harbor clinically relevant immune and molecular alterations. Prior studies by our group and others have demonstrated associations between immune features in tumor-adjacent lung tissue and patient outcomes, including survival and patterns of immune infiltration (13,14). Importantly, tumor-adjacent tissues exhibit inflammatory, immune, and molecular alterations that may reflect early microenvironmental remodeling or local effects of the tumor (13,15–17). Despite these observations, the spatial organization of immune cells within the tumor-adjacent tissue remains less well characterized than that of tumor tissue, particularly in matched tissue specimens from patients with early-stage NSCLC.

In this study, we leveraged a cohort of predominantly early-stage NSCLC tumor tissues and matched tumor-adjacent uninvolved lung tissues stained using multiplex immunofluorescence (mIF), spectrally imaged through Vectra, and consolidated into comprehensive immunoprofiling data files. Here, we characterize spatial differences between tumor and tumor-adjacent uninvolved lung tissues in early-stage NSCLC and evaluate associations between tissue-specific spatial features and clinical characteristics.

## 2 Methods

### 2.1 Cohort and Study Design

This study utilized multiplex immunofluorescence (mIF) data and whole-exome sequencing (WES) data from the UT MD Anderson Cancer Center ICON (ImmunogenomiC PrOfiling of Non-Small Cell Lung Cancer) cohort of NSCLC tumor and adjacent uninvolved lung tissues (PA15-111). Histology groups other than lung adenocarcinoma and lung squamous cell carcinoma were excluded from analysis. A total of 101 patient tumor samples were included, of which 91 (90%) had matched tumor-adjacent uninvolved samples. The protocol for multiplex immunofluorescence and digital image acquisition was previously validated, described and published with the aim of characterizing epithelial cells (malignant cells or epithelial cells in adjacent tumor tissue) and different immune cells within the tumor microenvironment and adjacent tissue (18–21). Specifically, we used the following panels: Panel 1 (**P1: PD-L1/PD-1 & TIL**) included biomarkers to identify epithelial cells (CK), T cells (CD3 and CD8) and macrophages (CD68), while also assessing PD-1 and PD-L1 immune checkpoints in epithelial and immune cells. Panel 2.1 (**P2.1: B-cell & NK cell activation**) targeted epithelial cells (CK), B-cells (CD20), and Natural Killer (NK) cells (CD57), while assessing functional activation (Granzyme B) and memory markers (CD45RO) in immune cells. Panel 2.2 (**P2.2: T-cell Activation**) identified epithelial cells (CK) and T cells (CD3), including cytotoxic (CD8) and regulatory (FOXP3) subsets, while assessing functional markers to identify memory T cells (CD45RO) and activated T cells (Granzyme B) (18–21). Single cells were plotted at their segmented coordinates over the mIF composite image for each region of interest (ROI) and colored by harmonized phenotype, using each antibody’s acquisition color (single-marker phenotypes) or the blend of those colors (multi-marker phenotypes); marker-negative and unclassified cells were shown in grey. Overlays were generated in R with ggplot2.

### 2.2 Spatial Analysis

Comprehensive immunoprofiling files were processed using a custom R pipeline built around the SPIAT package (22). Cell coordinates, phenotype annotations, and normalized marker intensities were extracted for each ROI. The ROIs were predefined by the source dataset and included as provided, with no study-specific ROI selection or exclusion. Normalized marker intensities were utilized for quality control, whereas cell coordinates and phenotypes were used for spatial analyses. Cells were further classified into broad lineage groups distinguishing tumor and epithelial cells (CK^+^), T cells (CD3^+^), Cytotoxic T cell subsets (CD3^+^CD8^+^), macrophages (CD68^+^), NK-like cells (CD57^+^), B cells (CD20^+^), memory T cells (CD45RO^+^), and regulatory cells (FOXP3^+^).

Cell densities (cells/mm^2^) were calculated for each lineage group and phenotype. ROI area was estimated from the minimum and maximum X and Y coordinates, with the resulting rectangular area expressed in pixels. Using a scale factor of 0.5 microns per pixel at 20x magnification, areas were converted to square microns and then to square millimeters by dividing by 10^6^. The following analyses were performed at the cell-phenotype level.

Spatial heterogeneity was quantified using SPIAT’s Prevalence and Distinctiveness metrics. The Prevalence of a pattern is defined as the proportion of grid squares in which a pattern is present, whereas Distinctiveness reflects the overall frequency of that pattern within an image. Prevalence was calculated using localized entropy, which split an ROI into 5×5 fishnet grid squares based on a chosen entropy threshold (0.75) for a spatial pattern and calculating the percentage of grid squares, denoting the prevalence of a spatial pattern. Distinctiveness of a spatial pattern was calculated using Global Moran’s I, where a high distinctiveness score indicates clustering in the tissue, a distinctiveness score of zero indicates random distribution, and a low distinctiveness score indicates dispersion throughout the tissue (22).

Average Nearest Neighbor Index (ANNI), which evaluates the spatial clustering or dispersion of specific cell types, was calculated using SPIAT. As part of the SPIAT module, the z score and p-value of the ANNI was calculated to validate the significance of the spatial pattern, with a p-value threshold of 0.05. High ANNI scores (ANNI>1) indicate dispersion, low ANNI scores (ANNI<1) indicate clustering, and ANNI scores at approximately 1 (ANNI ∼ 1) indicate random distribution (22).

Cell colocalization, which evaluates the spatial clustering or dispersion of a pair of cell types, was calculated using SPIAT by computing the inhomogeneous Ripley’s L-cross function (a variance-normalized version of the Ripley’s K statistic) for all ordered cell-type pairs, then calculating the area under the curve (AUC) of the cross-L function. Negative AUC values indicate segregation or low colocalization, while positive AUC values indicate aggregation of cell types or high colocalization (22,23).

ROI features per patient and tissue type were aggregated to avoid treating ROIs as individual samples. Aggregation of the ROI densities was calculated through regular averaging, while aggregation of the spatial metrics was calculated through weighted averaging using the Landau average in the spagg R package (24).

Hierarchical neighborhood clustering was performed by creating an N-by-N consensus matrix of the mean aggregated L-cross AUC values for each cell-type pair. Hierarchical clustering was applied to the rows of this consensus matrix using Euclidean distance with Ward.D2 linkage, grouping cell types by similarity of their outgoing co-localization profiles. The optimal number of clusters (k=2) was selected by maximizing mean silhouette width, subject to a minimum cluster size of 3 cell types (cell types or patients) (25).

Colocalization networks were constructed from the aggregated L-cross AUC values, with nodes representing cell phenotypes and directed edges representing the mean AUC for the directional spatial relationship between sender and receiver cell types. Networks were generated separately by marker panel and tissue compartment, excluding homotypic pairs. Each directed pair was tested against a null hypothesis of 0, corresponding to complete spatial randomness, using a one-sample Wilcoxon signed-rank test; analyses required at least 10 patients per stratum and at least 5 contributing patients per edge, with Benjamini-Hochberg correction applied across all pairs within each stratum. Edges were retained at adjusted p≤0.05, with edge color indicating attraction versus avoidance, edge width indicating mean AUC, and node size reflecting the number of significant incident edges. To permit direct visual comparison, all panels shared a common node set comprising the union of cell phenotypes present in either compartment, a fixed circular layout, and common edge-width and node-size scales; nodes with no significant edges in a given compartment were retained and displayed at reduced opacity (26,27).

### 2.3 Statistical Analysis

Associations between cellular features, spatial interaction measures, or cluster assignments and clinical, pathological, or genomic variables were assessed using nonparametric and correlation-based methods. Two-group comparisons were performed using Wilcoxon rank-sum tests, while multi-group comparisons were performed using Kruskal-Wallis tests. Associations between quantitative variables were evaluated using Spearman-rank correlation, and relationships between categorical variables in cluster-based analyses were tested using Fisher’s exact test. Effect sizes were reported as Spearman’s rho, rank-biserial correlation, or epsilon-squared, corresponding to the continuous, binary, and categorical comparisons, respectively. Tumor versus normal comparisons were conducted using Wilcoxon signed-rank tests for matched samples.

Survival outcomes, including overall survival and recurrence-free survival, were analyzed using Kaplan-Meir methods with log-rank tests and Cox proportional hazards regression models. Predictors were evaluated as continuous variables, median-dichotomized variables, and as the difference between tumor and tumor-adjacent values (Δ = tumor value – tumor-adjacent value) for metrics measured in both compartments. Results are reported as hazard ratio (HR) with 95% confidence intervals

Across all analyses, multiple testing was controlled by using the Benjamini-Hochberg false discovery rate method, and statistical significance was defined as q≤0.05. Minimum sample size thresholds were applied to ensure robustness of statistical testing.

## 3 Results

### 3.1 Tumor tissues exhibit higher immune cell densities than their adjacent counterparts

We utilized mIF ROI coordinates from 101 tumor samples and 91 matched adjacent normal samples from the ICON cohort to perform spatial analysis (**Figure 1A**) utilizing three mIF panels to characterize the tumor microenvironment: (1) PD-L1/PD-1 and tumor-infiltrating lymphocytes (TILs); (2) B-cell and NK-cell activation; and (3) T-cell activation (**Figure 1B**). To validate cell phenotypes, we overlaid X-Y coordinates onto representative ROIs. Phenotyped cells aligned with underlying staining patterns, supporting the accuracy of both cell identification and spatial localization, supporting subsequent spatial analyses (**Supplementary Figures 1-3**).

**Figure 1.**
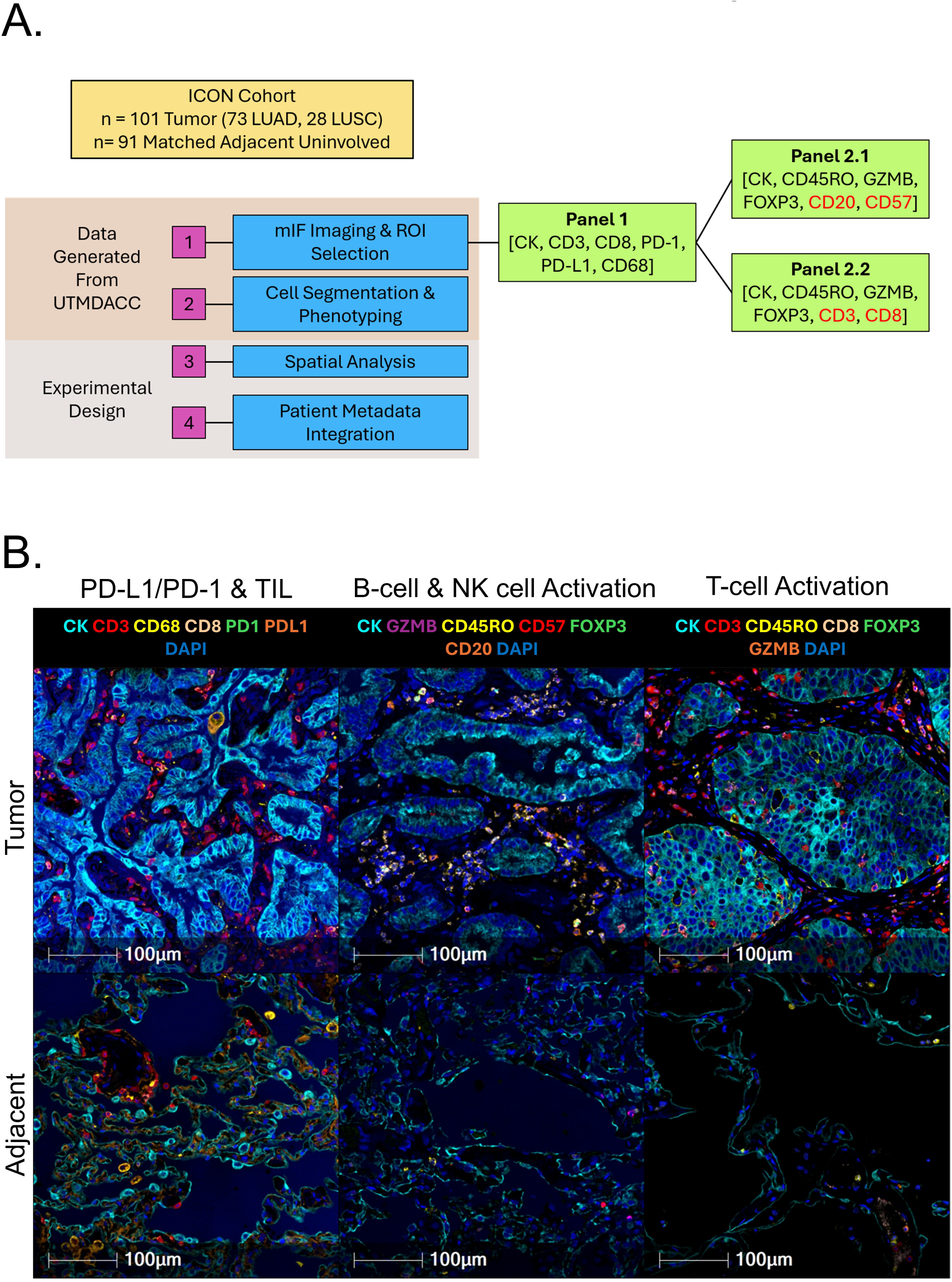
Patient cohort and multiplex immunofluorescence (mIF) data overview. **(A)** Overview of the ICON cohort analyzed, providing brief details on data generation and spatial analysis. **(B)** Representative mIF images of tumor and matched adjacent normal tissues for panels 1, 2.1, and 2.2 with corresponding cell-type annotations.

We first evaluated the cellular composition across the cohort utilizing density (cells/mm^2^) with all panels displaying elevated densities for most immune cells in tumor tissue compared to tumor-adjacent tissue for both cell lineage (**Figure 2A**) and phenotypic markers (**Supplementary Figure 4A**). Specifically, tumor samples displayed significantly higher immune cell densities, both at the lineage and specific phenotype levels, compared to their matched tumor-adjacent tissues for T cells, macrophages, B cells, and memory cells (p<0.001, **Figure 2B-F, Supplementary Figure 4B-F**).

**Figure 2.**
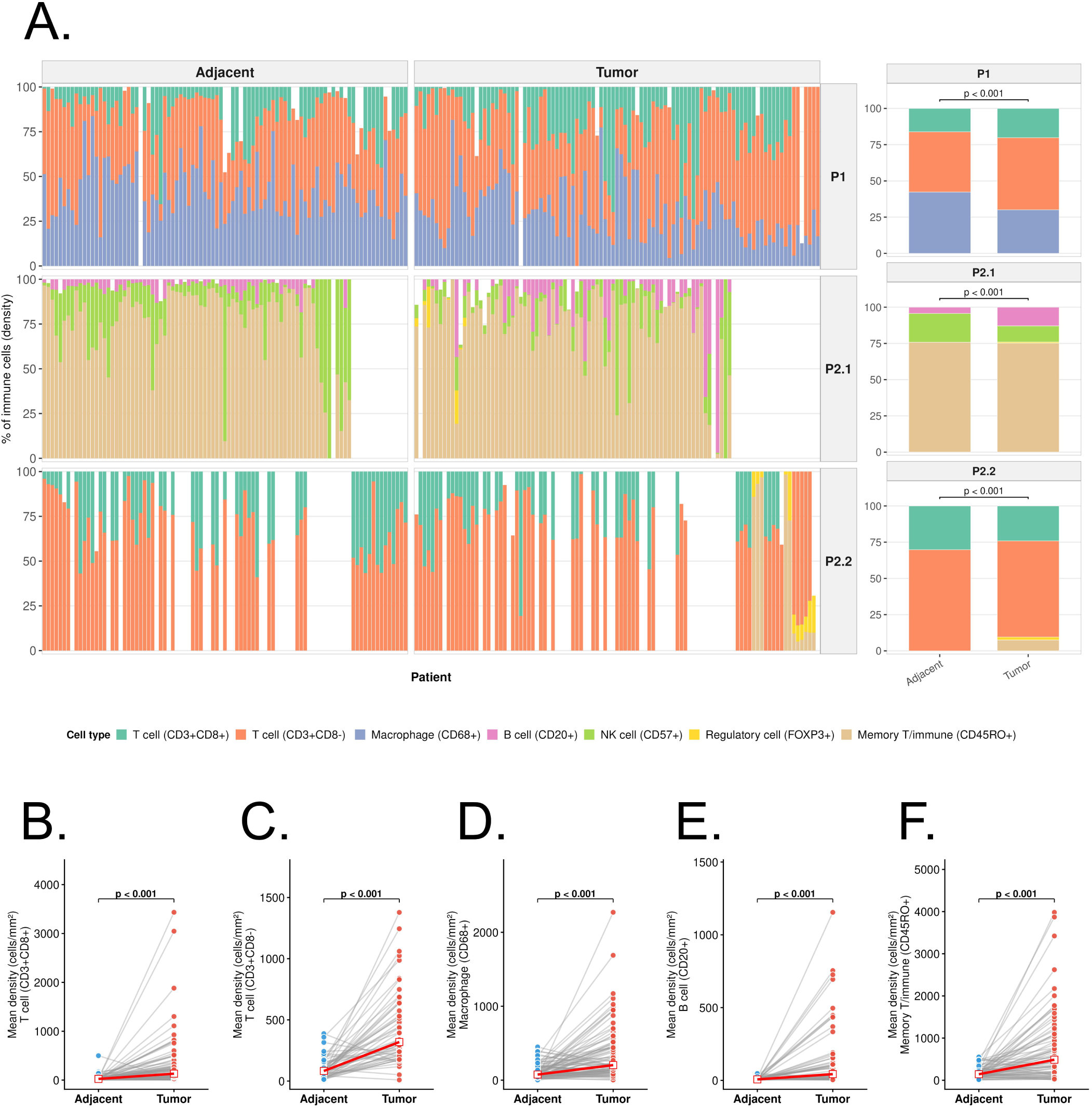
Most immune cell densities are elevated in tumor samples compared to adjacent uninvolved samples. **(A)** Lineage cell type densities (cells/mm^2^) show an increase in T cells, B cells, and regulatory cells in tumor samples, while adjacent samples show increased densities of macrophages and natural killer (NK) cells. **(B-F)** Pairwise comparisons of T cells (B, C), macrophages (D), B cells (E), and memory T/immune cells (F) show that these immune cell densities are elevated in tumor samples compared to adjacent tissues.

### 3.2 Tumor tissues exhibit higher spatial heterogeneity and localized organization than tumor-adjacent tissue

We next evaluated spatial heterogeneity across the cohort, calculating Prevalence and Distinctiveness of a spatial pattern. The Prevalence of a pattern is defined as the proportion of grid squares in which a pattern is present, whereas Distinctiveness reflects the dispersion or localization of a pattern based on spatial autocorrelation (measured by Moran’s I). Across the cohort, tumor samples exhibited a wide range of Prevalence values (0-100%), indicating substantial inter-patient variability in spatial pattern prevalence and high spatial heterogeneity (**Figure 3A**). On average, tumor samples showed higher Prevalence of a spatial pattern compared to tumor-adjacent tissues (47.67% vs 31.21%, p<0.001). Tumor samples had a mean Distinctiveness score of 0.10 with most tumor samples exhibiting scores consistent with spatial patterns concentrated in localized regions and reflecting substantial spatial heterogeneity across tumors (**Figure 3B**).

**Figure 3.**
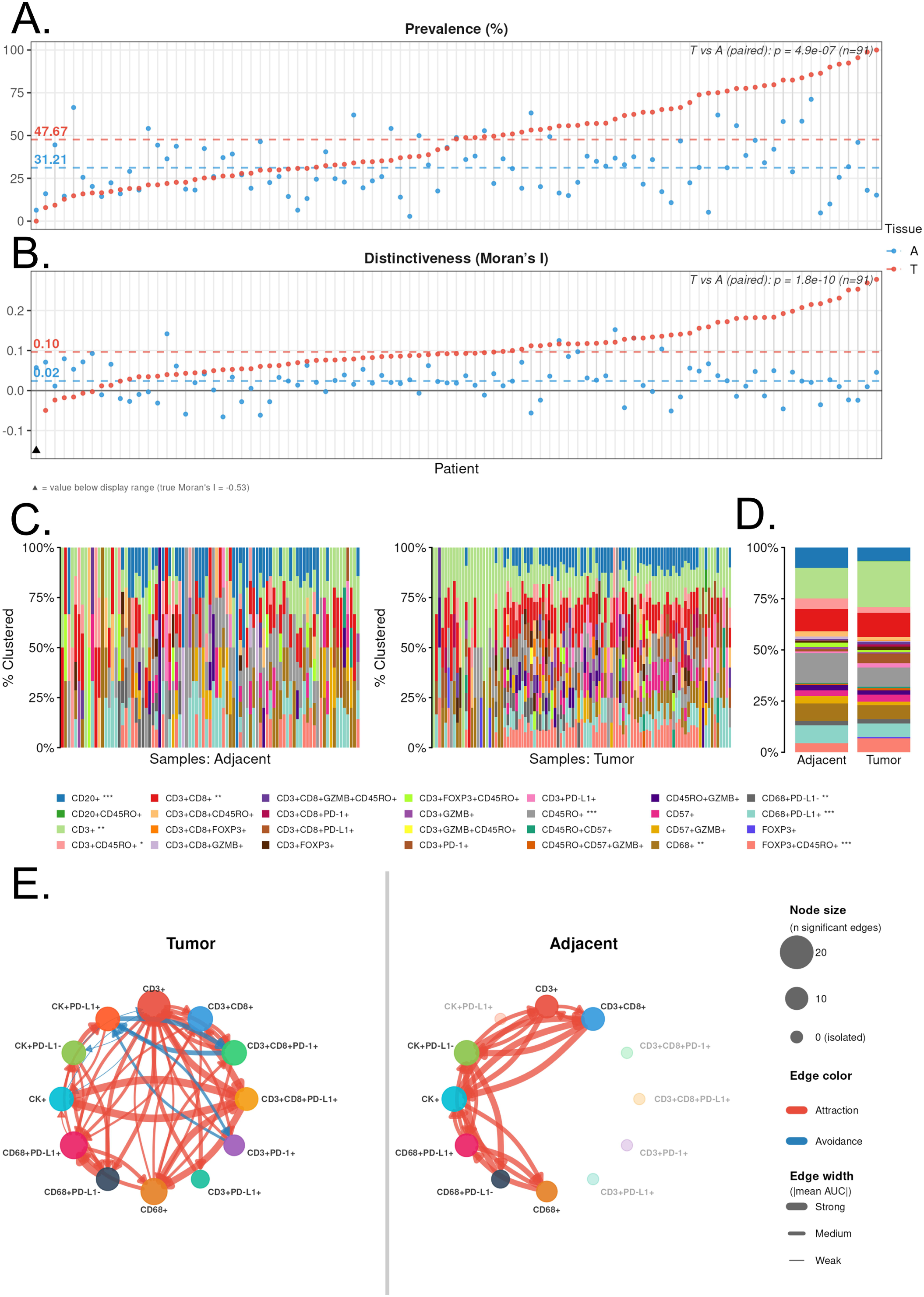
Tumor samples have distinct spatial architecture and interactions compared to adjacent uninvolved tissues. **(A)** Prevalence across patients ordered by increasing tumor value along the x-axis for (A) and (B). **(B)** Distinctiveness (global Moran’s I), reflecting the frequency/clustering of a pattern, per patient for tumor (red) and adjacent (blue) samples. Most tumor samples show positive spatial autocorrelation (Moran’s I > 0, indicating clustering), whereas adjacent samples are distributed around zero, including negative values (dispersion). Mean Moran’s I is 0.10 in tumor vs. 0.02 in adjacent tissue (p<0.001). Dashed horizontal lines indicate group means. One adjacent tissue point falls below the plotted range (true value = −0.53), marked with ▲. **(C)** Individual cell clustering across adjacent normal and tumor tissues reveals distinct distribution patterns among immune cell phenotypes: tumor samples display relatively consistent clustering profiles across samples, with similar clustering proportions of CD3+ T-cells, CD45RO+ subsets, and CD68+PD-L1+ cells, whereas adjacent tissues exhibit greater inter-sample variability in cell clustering across many immune cell phenotypes. **(D)** Averaged individual cell clustering in tumor-adjacent and tumor samples, showing greater percentages of most cells in tumor-adjacent compared to tumor tissue; asterisks are displayed in the shared legend next to cell types with statistically significant differences between the averaged tumor and tumor-adjacent values (*: p <0.05, **: p<0.01, ***: p<0.001). **(E)** Colocalization networks of L-cross AUC values for Panel 1, shown separately for tumor (left) and adjacent (right) tissue. Each node represents a cell type, and each arrow a directional spatial relationship from the reference to the target phenotype. Node size reflects the number of significant relationships involving that cell type; faded nodes had no significant relationships in that compartment. Edges connect phenotypes whose mean AUC differed significantly from spatial randomness across patients (one-sample Wilcoxon signed-rank test, BH-adjusted q ≤ 0.05), with edge color denoting the direction of association (red = attraction, blue = avoidance) and edge thickness scaling with |mean AUC|. Both panels share a common node set, layout, and scales; homotypic pairs were excluded.

Conversely, tumor-adjacent samples displayed consistently lower Prevalence and a narrower range (∼0-70%) compared to tumor samples, indicating that a pattern was present in fewer regions of the tissue. Distinctiveness values in tumor-adjacent tissues spanned both negative and positive Moran’s I values, reflecting lower apparent spatial heterogeneity and greater variability in spatial organization, from dispersed to confined patterns. Overall, within the sampled regions, tumor tissues showed higher spatial heterogeneity with high prevalence of a spatial pattern, and more consistently localized spatial patterning than adjacent tissues, whereas tumor-adjacent tissues showed lower prevalence of a spatial pattern compared to tumor, but with non-localized spatial organization. For 13 samples in which adjacent tissues exhibited higher Moran’s I than tumors, analysis revealed that the corresponding ROIs of tumor samples were predominantly composed of malignant cells (median of 76.2%), with limited representation of other cell types relative to the matched adjacent ROIs.

### 3.3 Distinct cell clustering and composition patterns appear in tumor versus tumor-adjacent tissues

To further characterize cell-level spatial organization, we evaluated the Average Nearest Neighbor Index (ANNI) in tumor and tumor-adjacent tissues, which quantifies whether specific cell types are clustered, randomly distributed, or more regularly spaced. We compared the clustered and random distributions of individual cell types across both tissue types, which revealed clear differences in tissue organization (**Figure 3C-D**, **Supplementary Figure 5A**).

Both tissue types contained individual cell clustering and random distributions, yet tumor-adjacent tissues showed a higher proportion of clustering for several individual cell populations, indicating preserved tissue architecture and baseline immune compartmentalization relative to tumor tissue. Cluster proportions were also heterogeneous across samples, with dominant clustered states observed in T-cell subsets, memory cell subsets, and macrophage subsets.

In contrast, tumor samples displayed lower proportions for some cell populations, consistent with a redistributed but population-specific spatial pattern. Among cell states with clustered spatial organization, tumor samples showed a trend toward more uniform proportions across patients than tumor-adjacent samples (median coefficient of variation: 0.39 vs 0.44, respectively), though the difference was not statistically significant (p=0.12). Together, these findings indicate that tumor tissues exhibit reduced individual cell clustering compared to their matched adjacent tissues, but more consistent cell-state composition across patients, whereas adjacent tissues display greater variability of composition across patients and stronger clustering for several individual cell populations.

### 3.4 Tumor tissues exhibit more selective immune spatial structure than tumor-adjacent tissues

We next evaluated spatial interactions between immune cell populations using area under the curve (AUC) values derived from Ripley’s L-cross function in tumor and tumor-adjacent samples. Unlike our previous analyses, which quantified the spatial organization (clustering or random distribution) of individual immune cell populations, Ripley’s L-cross specifically measures the spatial relationship between two distinct cell types, allowing us to assess whether particular immune subsets preferentially colocalize or spatially segregate. AUC, or colocalization values summarize these interactions across the range of distances examined, providing an integrated measure of pairwise spatial association.

Across immune cell pairs, tumor samples generally exhibited positive median AUC values, indicating a tendency for immune cell subsets to appear in close proximity (**Supplementary Figure 5B**). These findings reveal a spectrum of spatial relationships among immune cell populations in tumors, ranging from widespread colocalization to selective spatial segregation between specific cell-type pairs.

Among the pairs with negative colocalization values in tumor samples, some differed significantly compared to tumor-adjacent samples, including: (1) effector/memory cytotoxic T cells (CD3^+^CD8^+^CD45RO^+^) ↔ activated cytotoxic T cells (CD3^+^CD8^+^GZMB^+^) (p.adj<0.05); and (2) effector/memory cytotoxic T cells (CD3^+^CD8^+^CD45RO^+^) ↔ regulatory T cells (CD3^+^FOXP3^+^) (p.adj<0.05). These findings suggest there is selective spatial segregation in tumor samples between B-cells and memory/regulatory cells, as well as among functionally distinct T-cell subsets. In contrast, tumor-adjacent samples displayed a more variable colocalization distribution, with both positive and negative medians representing a heterogeneous mixture of colocalized and dispersed immune cell pairs. For immune cells in proximity to CK^+^ cells, tumor-adjacent tissues had higher AUC values than tumor tissues (**Supplementary Figure 5C**). This is suggestive of greater immune cell proximity to epithelial cells in tumor-adjacent tissue and reduced immune cell proximity to malignant cells in tumor regions. Overall, these findings suggest that tumor tissues exhibit more selective pairwise immune spatial structure, whereas tumor-adjacent tissues display greater variability and strong immune-epithelial associations.

### 3.5 Hierarchical clustering and network analysis reveal distinct higher-order organization in tumor and tumor-adjacent tissues

To move beyond pairwise spatial associations and define higher-order organization, we performed hierarchical neighborhood clustering and correlation network analysis using colocalization values for immune cell and CK^+^ cell phenotypes across all three mIF panels. Hierarchical clustering consistently identified two major groups corresponding to higher and lower levels of colocalization (**Supplementary Figure 6A-B**). Tumor-adjacent tissues exhibited more distinct separation between these groups than tumor tissues, suggesting greater spatial segregation among cell-type interactions.

Colocalization network analysis revealed distinct organizational patterns across tissue compartments. In the PD-L1/PD-1 and TIL panel, tumor tissues were characterized by extensive immune-immune cell colocalization and several segregation relationships involving malignant cells (**Figure 3E**), whereas tumor-adjacent tissues showed immune-epithelial cell colocalization and lacked significant segregation relationships. Similar patterns were observed in the T-cell activation panel, where tumor tissues exhibited greater colocalization among T-cell subsets, while adjacent tissues showed stronger associations between immune and epithelial cells (**Supplementary Figure 6D**). In contrast, the B-cell and NK-cell activation panel demonstrated more extensive colocalization and segregation relationships in tumor-adjacent tissue than in tumor tissue (**Supplementary Figure 6C**), suggesting that patterns of B-cell and NK-cell organization differ between tissue compartments.

Overall, these analyses reveal distinct higher-order spatial organization between tumor and tumor-adjacent tissues. Tumors were generally characterized by stronger immune cell-immune cell associations, whereas tumor-adjacent tissues more frequently exhibited immune-epithelial cell associations, highlighting compartment-specific patterns of cellular organization.

### 3.6 Tumor and tumor-adjacent tissues exhibit spatial features associated with outcomes

Lastly, we evaluated associations between spatial features, genomic features, and clinicopathological data across the cohort and found that tumor tissues showed several statistically significant associations (**Figure 4A**). As anticipated, low neoantigen burden was associated with decreased CK^+^ tumor cell density, however, high neoantigen burden was associated with increased cell densities of CK^+^PD-L1^+^, CD3^+^FOXP3^+^ cells, and CD3^+^ cells potentially suggesting increased tumor immunogenicity.

**Figure 4.**
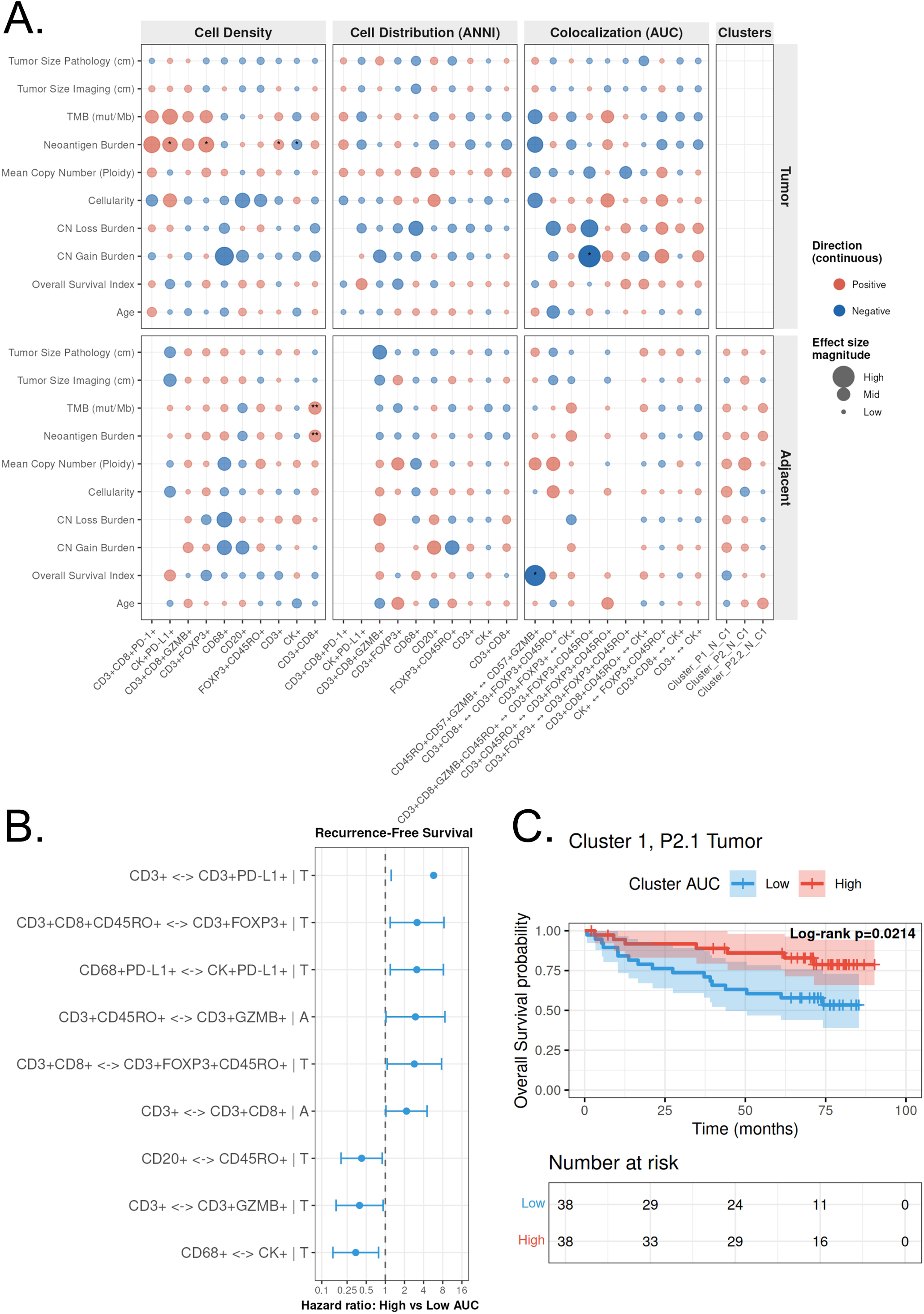
Associations between spatial metrics, clinical variables, and patient survival. **(A)** Bubble plot illustrates associations between patient genomic data, clinicopathological data, and spatial metrics, including cell density, average nearest neighbor index (ANNI), Lcross AUC, and hierarchical clustering results for tumor and adjacent uninvolved tissues. Bubble size represents effect size, while color indicates the direction of the association (red = positive; blue = negative). **(B)** Forest plot of hazard ratios for recurrence-free survival associated with bidirectional cell–cell colocalization pairs showing nominal significance (unadjusted p<0.05). High colocalization (high AUC) was associated with improved recurrence-free survival for interactions with hazard ratios < 1 and increased recurrence risk for interactions with hazard ratios > 1. None of these associations remained significant after multiple-testing correction. **(C)** Kaplan–Meier overall survival analysis of tumor samples stratified by high versus low L-cross AUC for Cluster 1 of Panel 2.1. Patients with high AUC values demonstrated significantly improved overall survival compared with those with low AUC values (log-rank p = 0.0214).

Copy number (CN) gain burden was negatively associated with increased spatial colocalization (L-cross AUC) between CD3^+^CD8^+^GZMB^+^CD45RO^+^ T-cells and CD3^+^FOXP3^+^CD45RO^+^ T-cells. In tumor-adjacent tissue, increased CD3^+^CD8^+^ T-cell density was associated with TMB (mut/Mb) and neoantigen burden, and increased colocalization between CD45RO^+^CD57^+^GZMB^+^ cells and CD57^+^GZMB^+^ cells was negatively associated with overall survival index.

Beyond these statistically significant findings, several observations emerged. In tumor tissue, neoantigen burden and TMB (mut/Mb) were trending positively with CK^+^PD-L1^+^, CD3^+^CD8^+^GZMB^+^, CD3^+^FOXP3^+^, and CD3^+^ densities. This is consistent with the expected relationship between these metrics, as neoantigens can arise from nonsynonymous somatic mutations, and tumors with a higher mutational burden tend to have a larger neoantigen repertoire (28). In tumor-adjacent tissue, the same positive trends with tumor-derived neoantigen burden and TMB (mut/Mb) were observed across immune cell densities, suggesting that the influence of tumor mutational and neoantigen burden on immune cell infiltration may extend into the surrounding tissue microenvironment.

Although few associations were observed with colocalization against genomic and clinicopathological data, several cell-pair colocalization metrics showed nominal associations with recurrence-free survival (RFS) prior to multiple-testing correction (**Figure 4B**). Broadly, increased colocalization between immunosuppressive and effector immune populations was associated with worsened RFS in both tumor and tumor-adjacent tissues, whereas improved RFS was associated with increased colocalization among effector, memory, and B-cell subsets in tumor tissue. The consistency of these associations across both directions of each cell-pair suggests that they may reflect genuine spatial relationships.

Although no genomic or clinicopathological data associations with hierarchical clustering across the cohort met the predefined significance threshold of p<0.05 (**Figure 4A**), increased cluster colocalization in the B-cell and NK-cell activation panel in tumor tissue was associated with improved recurrence-free survival, while low cluster colocalization was associated with worsened recurrence-free survival (p<0.05, **Figure 4C**). Taken together, these findings highlight that both tumor and adjacent tissues may harbor distinct, spatial signatures associated with clinicopathological features, genomic features, and patient outcomes.

## 4 Discussion

In many NSCLC studies, tumor-adjacent “normal” tissues are used primarily as a baseline comparison for tumor samples (29–32). However, we and others have previously shown that tumor-adjacent tissues in NSCLC that are immune-active or inflamed can correlate with patient outcomes (33,13,14). Although one study found that adjacent tissues alone are insufficient for making associations with clinical outcomes, these findings suggest that adjacent tissues may still provide additional biological and spatial insights that warrant further investigation (15). Previous spatial studies involving adjacent tissues in NSCLC utilized few visualization markers, while spatial transcriptomic studies have focused on tumor tissue rather than the prognostic value of the adjacent tissues (34–36). Thus, the spatial organization and prognostic significance of tumor-adjacent tissues remain poorly defined. Here, we performed spatial analysis on multiplex immunofluorescence data obtained from 101 tumor and 91 matched tumor-adjacent samples from early-stage NSCLC patients.

Despite greater compositional heterogeneity in tumors, the sampled tumor regions showed more recurrent and localized cell patterning than tumor-adjacent tissue. This suggests that heterogeneity within the tumor may be self-organized into nonrandom micro-niches, whereas adjacent tissue showed consistently broader defined spatial relationships (34,37,38).

Analysis of individual cell distribution highlighted that tumor tissue is characterized by a shift in spatial organization from stronger local clustering in adjacent tissue to a more redistributed but patient-consistent cellular architecture in tumor regions, suggesting a change in tissue organization rather than a simple shift in clustering (39).

Colocalization results suggest that tumor tissues exhibit more conserved and selective pairwise immune spatial structure, whereas tumor-adjacent tissues display greater variability and strong immune-epithelial associations. This likely reflects the fact that the tumor microenvironment actively reorganizes local immune architecture, leading to more selective immune spatial relationships within tumor regions and broader, more variable interactions in adjacent tissue (40,41).

Hierarchical clustering and colocalization network analyses revealed distinct higher-order spatial organization between tumor and tumor-adjacent tissues. Tumor-adjacent tissues more frequently exhibited immune cell–epithelial cell associations and showed greater separation between highly and moderately colocalized cell groups, suggesting more spatially segregated cellular interactions. In contrast, tumors were generally characterized by stronger immune cell–immune cell associations. These patterns varied across mIF panels, indicating that the organization of the immune microenvironment is context- and cell-type dependent.

When comparing significant associations between spatial features and genomic data, we found that high tumor neoantigen burden was associated with increased density for CD3^+^ T-cells. This reflects the impact of neoantigen presentation on T-cell activation, resulting in greater TIL infiltration (42). Additionally, association with increased PD-L1^+^ tumor cells may represent a consequence of IFN-γ produced by neoantigen-reactive T-cells (43). Together, these findings are consistent with a neoantigen-driven immune response, in which enhanced T-cell recognition of tumor antigens is accompanied by adaptive PD-L1 upregulation on tumor cells.

Several spatial patterns showed potential relationships with recurrence-free survival, despite few significant associations being identified with genomic and clinicopathological data. Increased colocalization between immunosuppressive and effector immune populations, including regulatory and cytotoxic T-cell subsets, as well as between PD-L1^+^ macrophages and PD-L1^+^ malignant cells, may be linked to poorer recurrence-free survival.

Conversely, increased colocalization among effector, memory, and B-cell populations may be associated with more favorable outcomes (44,45). While these observations did not reach statistical significance, they suggest that the spatial organization of immune populations may influence antitumor immunity and clinical outcome, warranting further investigation.

Our study does exhibit certain limitations. First, immunofluorescence panels were restricted to five immune markers each, which limited the ability to comprehensively assess complex cellular states and interactions. Second, imaging was restricted to selected regions of interest (ROIs), which may not capture the full spatial heterogeneity of tumor and tumor-adjacent tissues and could possibly introduce sampling bias.

Overall, our study supports the concept that tumors are not simply disorganized and spatially heterogenous, but instead exhibit selectively structured spatial niches, while tumor-adjacent tissues retain a more variable and compartmentalized structure, and that spatial features of tumor and tumor-adjacent tissues may be associated with patient outcomes.

## 5 Data Availability

Datasets are available on request: Raw data supporting the conclusions of this article will be made available by the authors upon reasonable request.

All relevant statistical code will be deposited on GitHub upon acceptance.

## 6 Ethics Approval

This study utilized patient samples that were collected from NSCLC patients enrolled in the ICON study which began in April 2016 through September 2018. Informed consent was obtained from all study participants approved by the University of Texas MD Anderson Cancer Center’s Institutional Review Board.

## 7 Author Contributions

IP: Conceptualization, data curation, formal analysis, investigation, methodology, visualization, writing—original draft, writing—review and editing.

CA: Imaging, resources, writing–review and editing

JS: Writing–review and editing

AM: Writing–review and editing

EB: Writing–review and editing

PJ: Data curation, writing–review and editing

AP: Writing–review and editing

TK: Data curation, writing–review and editing

XW: Data curation, writing–review and editing

XS: Data curation, writing–review and editing

QL: Writing–review and editing

HI: Imaging, writing–review and editing

AT: Imaging, writing–review and editing

MJ: Imaging, writing–review and editing

CH: Writing–review and editing

AW: Writing–review and editing

AAV: Writing–review and editing

LK: Writing–review and editing

TC: Writing–review and editing

IW: Writing–review and editing

ICON Study Team: Data curation, imaging, writing–review and editing

LMS: Writing–review and editing

JVH: Writing–review and editing

JZ: Writing–review and editing

DLG: Writing–review and editing

AR: Conceptualization, resources, supervision, writing—original draft, project administration, writing—review and editing.

## 9 Acknowledgments

We thank the members of the ICON study team and Multiplex Immunofluorescence and Image Analysis Laboratory in the department of Translational Molecular Pathology at the University of Texas MD Anderson Cancer Center, for their contributions in image data acquisition and consolidation. The author(s) acknowledge the support of the High-Performance Computing for research facility at the University of Texas MD Anderson Cancer Center for providing computational resources that have contributed to the research results reported in this paper.

## 11 Generative AI statement

The author(s) declared that generative AI was not used in the creation of this manuscript.

---

**Supplementary Figure 1. Representative multiplex overlay and ROI-level validation of Panel 1 cell phenotypes. (A)** Representative overlay of a tumor ROI from Panel 1 (sample 13T) showing the original multiplex immunofluorescence image with phenotype-labeled cell coordinates derived from the raw segmentation files. **(B)** Representative overlay of a tumor-adjacent ROI from Panel 1 (sample 13N) showing the original multiplex immunofluorescence image with phenotype-labeled cell coordinates derived from the raw segmentation files.

**Supplementary Figure 2. Representative multiplex overlay and ROI-level validation of Panel 2.1 cell phenotypes. (A)** Representative overlay of a tumor ROI from Panel 2.1 (sample 3T) showing the original multiplex immunofluorescence image with phenotype-labeled cell coordinates derived from the raw segmentation files. **(B)** Representative overlay of a tumor-adjacent ROI from Panel 2.1 (sample 66N) showing the original multiplex immunofluorescence image with phenotype-labeled cell coordinates derived from the raw segmentation files.

**Supplementary Figure 3. Representative multiplex overlay and ROI-level validation of Panel 2.2 cell phenotypes. (A)** Representative overlay of a tumor ROI from Panel 2.2 (sample 118T) showing the original multiplex immunofluorescence image with phenotype-labeled cell coordinates derived from the raw segmentation files. **(B)** Representative overlay of a tumor-adjacent ROI from Panel 2.2 (sample 118N) showing the original multiplex immunofluorescence image with phenotype-labeled cell coordinates derived from the raw segmentation files.

**Supplementary Figure 4. Most immune cell phenotype densities are elevated in tumor samples compared to adjacent uninvolved samples. (A)** Specific immune cell phenotype densities (cells/mm^2^) show an increase in T cell, B cell, regulatory cell and memory immune cell subsets, while adjacent samples show increased densities of macrophages, natural killer (NK) cells, memory/immune cells, and T cell subsets. **(B-F)** Pairwise comparisons of specific phenotypes of CD3^+^ T cells (B), CD68^+^ macrophages (C), CD20^+^ B cells (D), CD3^+^CD8^+^ T cells (E), and memory/regulatory T (CD3^+^FOXP3^+^CD45RO^+^) cells (F) show that these immune cell phenotype densities are elevated in tumor samples compared to adjacent tissues.

**Supplementary Figure 5. Adjacent normal tissues also have altered spatial architecture and interactions when compared to tumor tissues. (A)** Random distribution of individual cells for adjacent normal tissues (left) and tumor tissues (right), showing similar percentages of most cells in tumor-adjacent compared to tumor tissue; asterisks are displayed in the shared legend next to cell types with statistically significant differences between the averaged tumor and tumor-adjacent values (*: p <0.05, **: p<0.01, ***: p<0.001). **(B)** In pairs seen in at least 60% of patients, colocalization analysis of immune cells shows that most cell pairs in tumor tissue have a positive AUC indicative of high colocalization, with adjacent tissues having a reduced AUC value compared to tumor tissues (*: p <0.05, **: p<0.01, ***: p<0.001). **(C)** In pairs seen in at least 60% of patients, colocalization analysis of immune cells and CK^+^ cells (epithelial cells for adjacent, and malignant cells for tumor) shows that adjacent tissues have a higher AUC than tumor tissues indicative of increased colocalization of immune cells to the epithelium compared to immune cells with malignant cells (*: p <0.05, **: p<0.01, ***: p<0.001).

**Supplementary Figure 6. Spatial colocalization analysis reveals distinct cell–cell interaction patterns between tumor and adjacent uninvolved lung tissue. (A, B)** Hierarchical clustering of pairwise colocalization AUC values identified two distinct clusters (k = 2) within each mIF panel for tumor (A) and adjacent uninvolved (B) tissues. Heatmaps display the mean AUC values for each cluster, with red indicating higher colocalization and blue indicating lower colocalization. Compared with tumor tissue, adjacent tissue exhibited a greater prevalence of lower AUC values, consistent with reduced spatial colocalization among immune cell populations. **(C, D)** Colocalization networks of L-cross AUC values for Panel 2.1 (C) and Panel 2.2 (D), shown separately for tumor (left) and adjacent (right) tissue. Each node represents a cell type, and each arrow a directional spatial relationship from the reference to the target phenotype. Node size reflects the number of significant relationships involving that cell type; faded nodes had no significant relationships in that compartment. Edges connect phenotypes whose mean AUC differed significantly from spatial randomness across patients (one-sample Wilcoxon signed-rank test, BH-adjusted q ≤ 0.05), with edge color denoting the direction of association (red = attraction, blue = avoidance) and edge thickness scaling with |mean AUC|. Both panels share a common node set, layout, and scales; homotypic pairs were excluded.

