## Supplementary figures and images for "Distinct Spatial Immune Architectures in Tumor and Tumor-Adjacent Tissues of Early-Stage Non-Small Cell Lung Cancer"

### Supplemental Figure 1

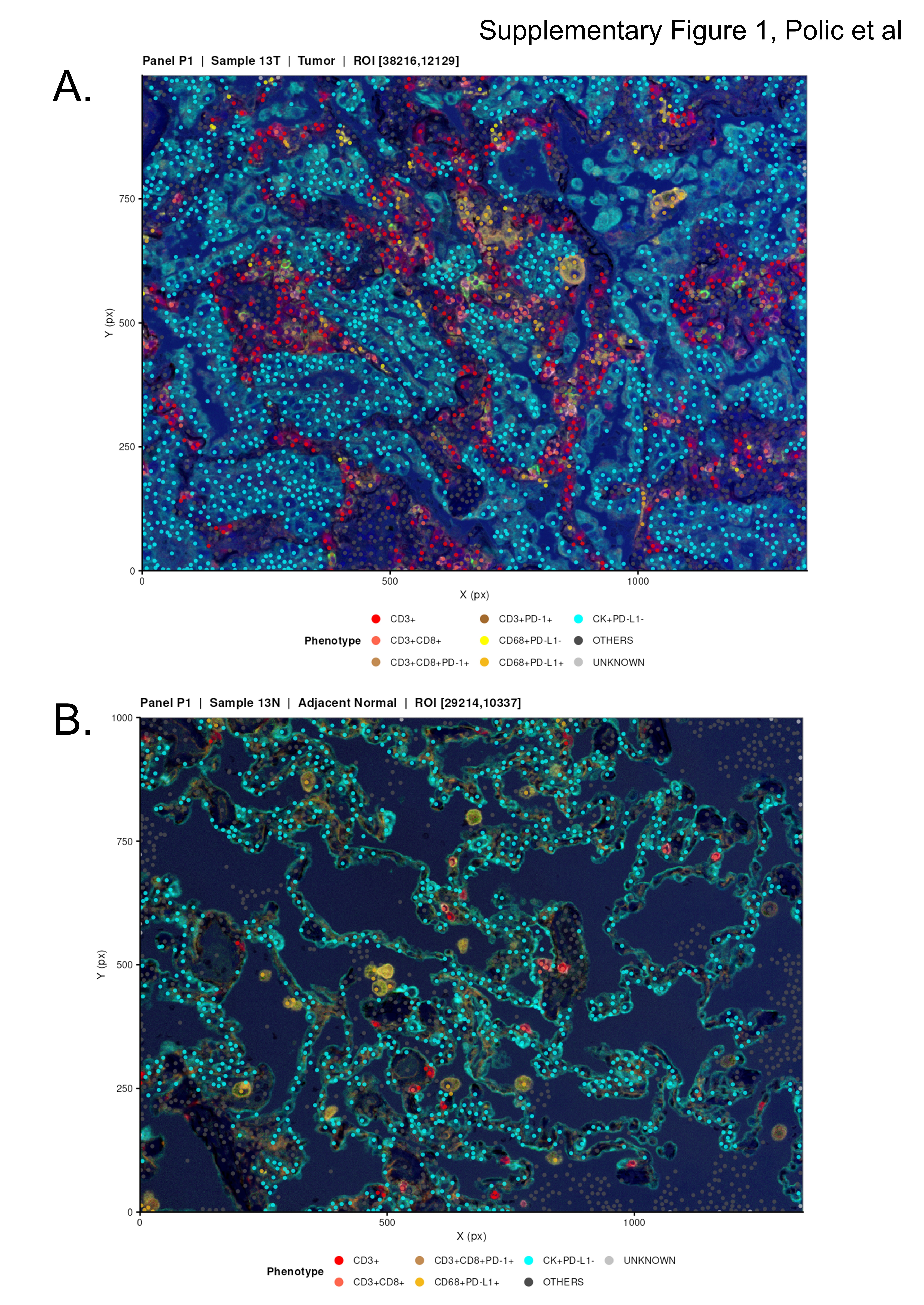

### Supplemental Figure 2

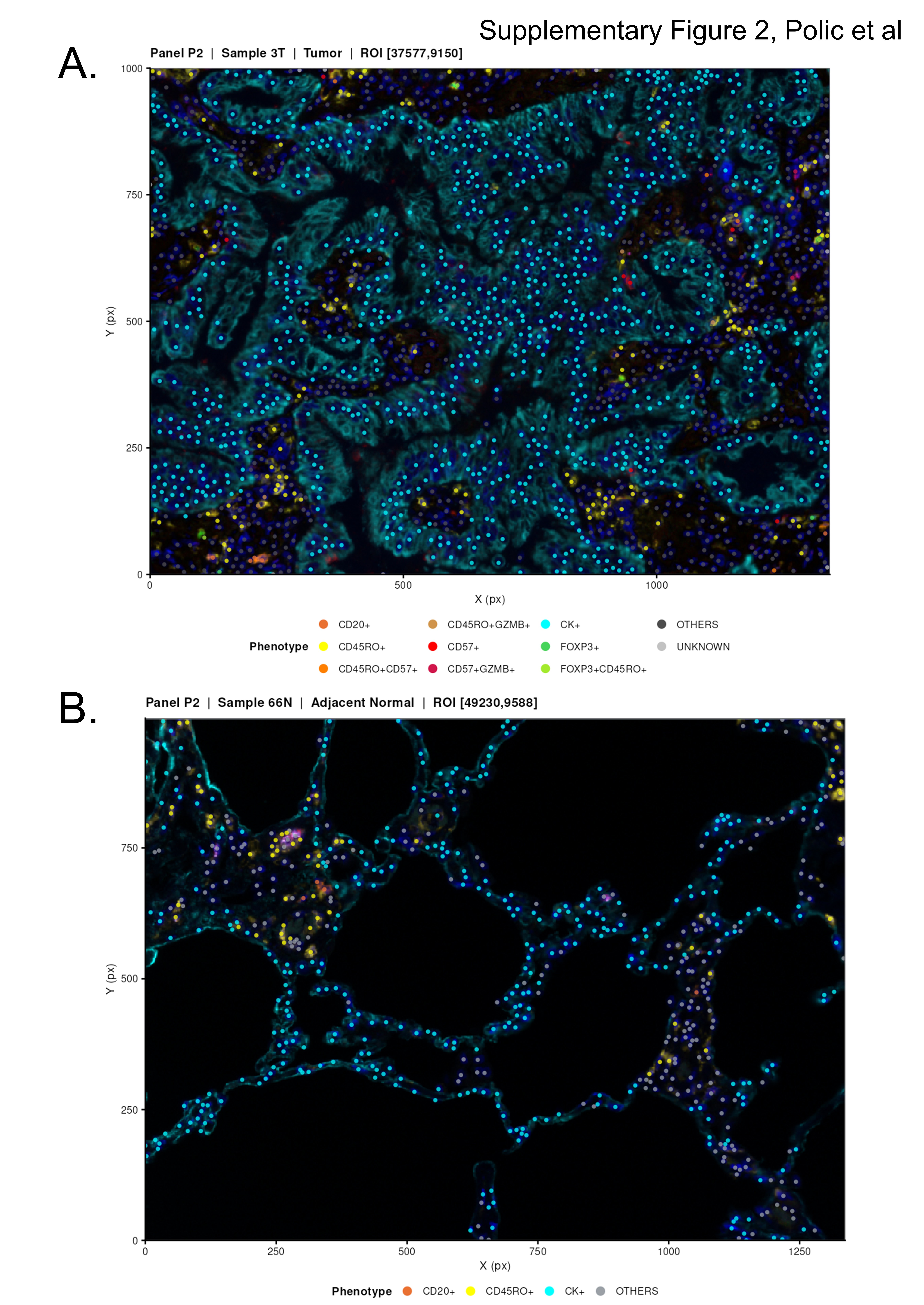

### Supplemental Figure 3

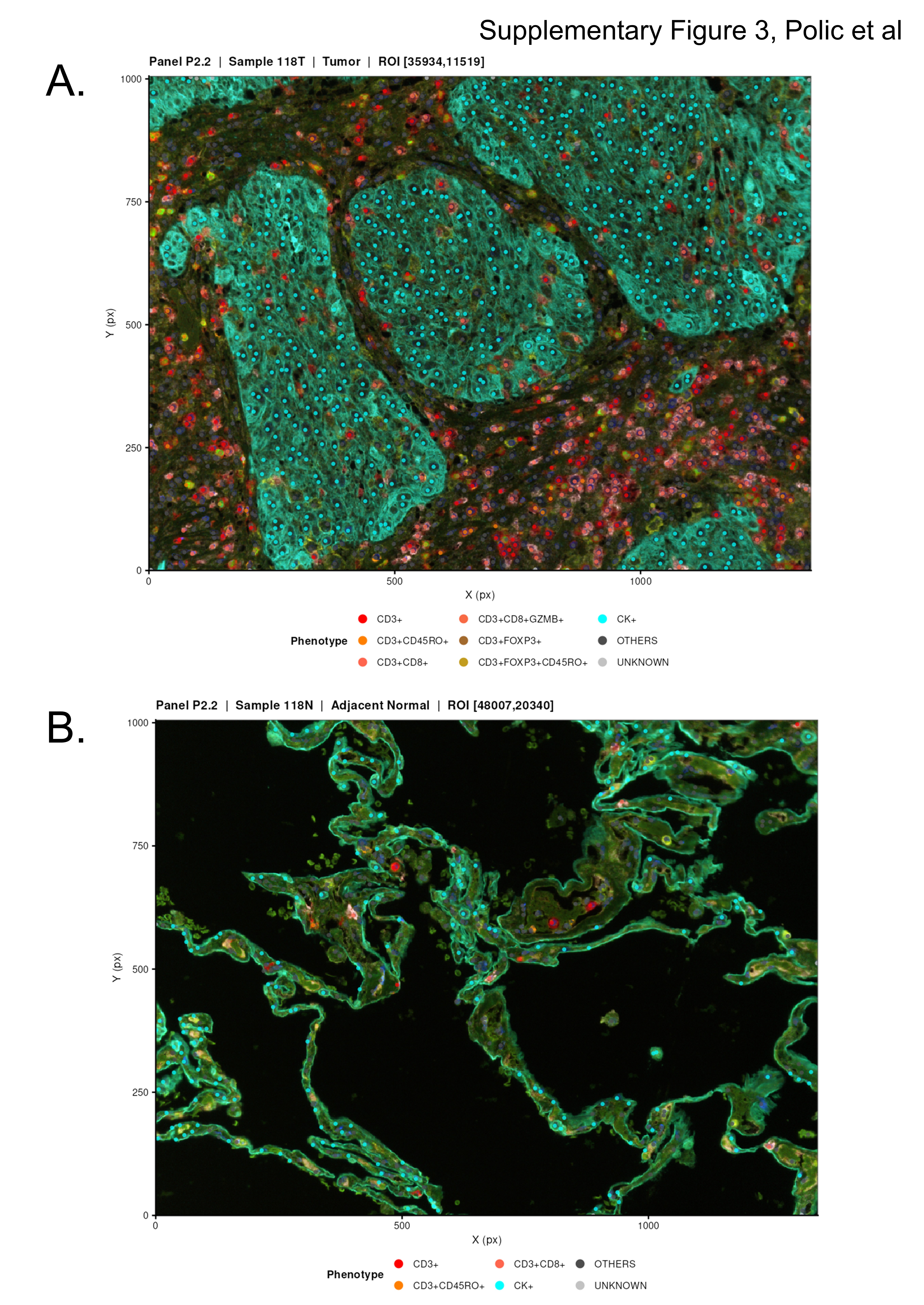

### Supplemental Figure 4

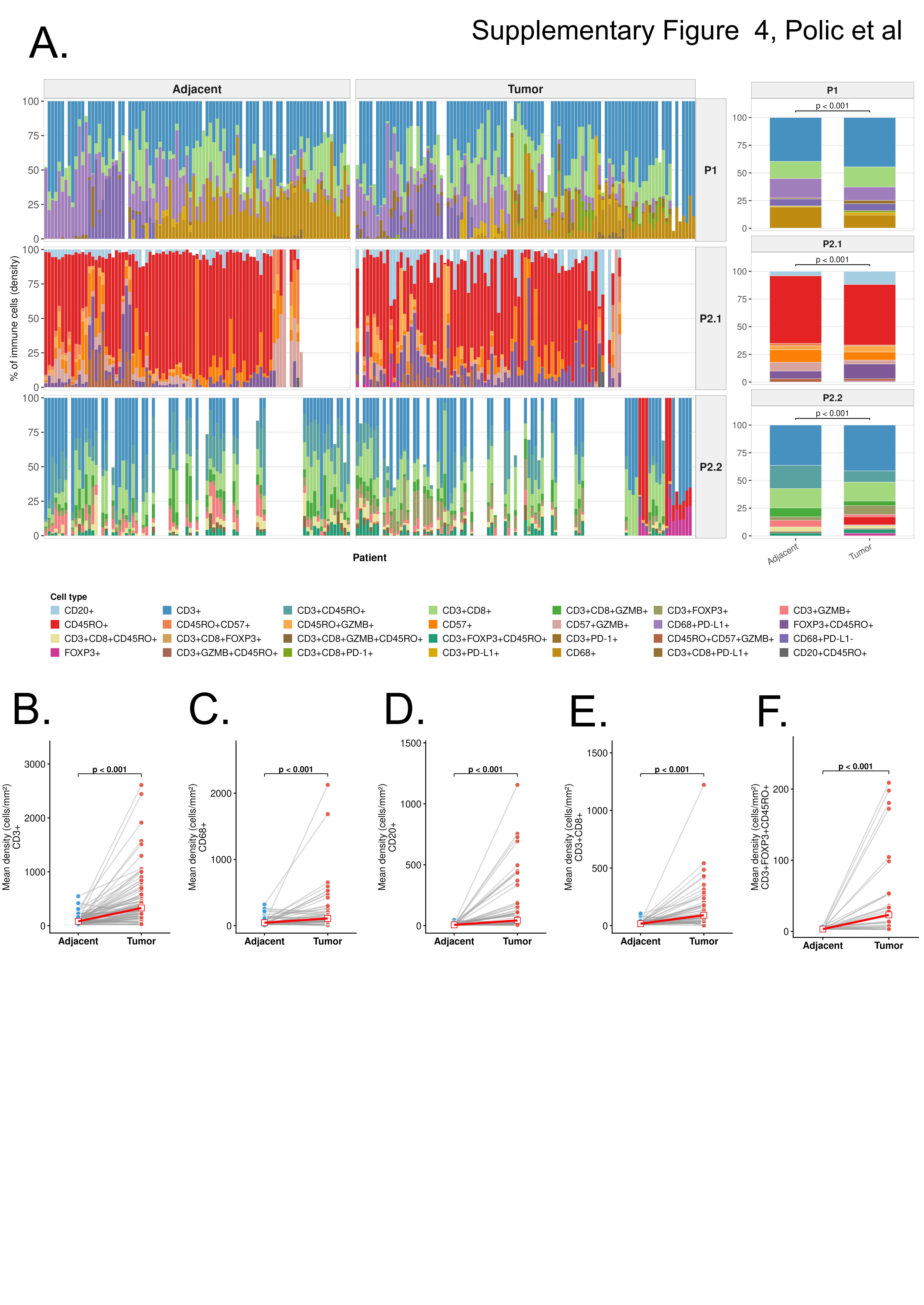

### Supplemental Figure 5

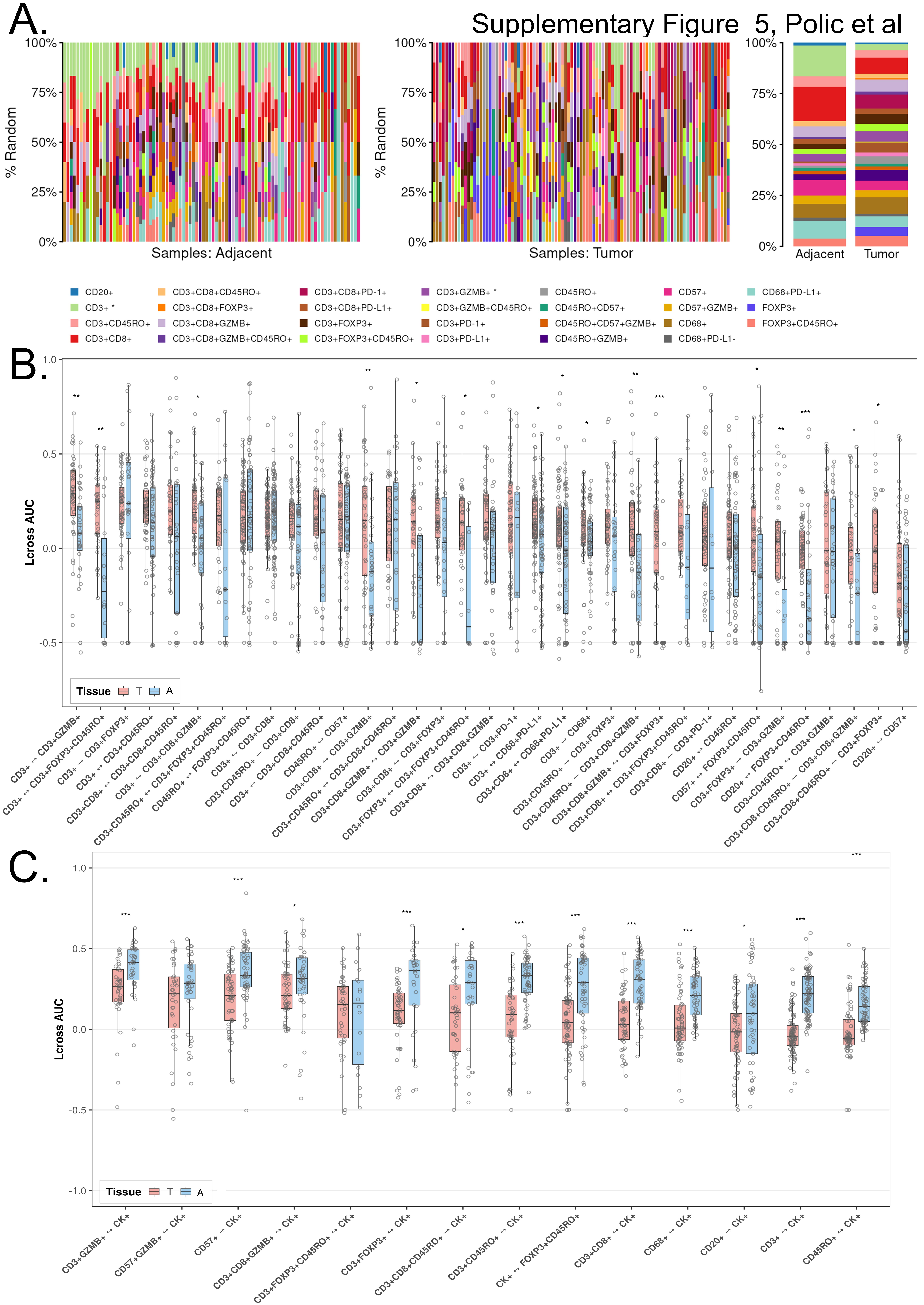

### Supplemental Figure 6

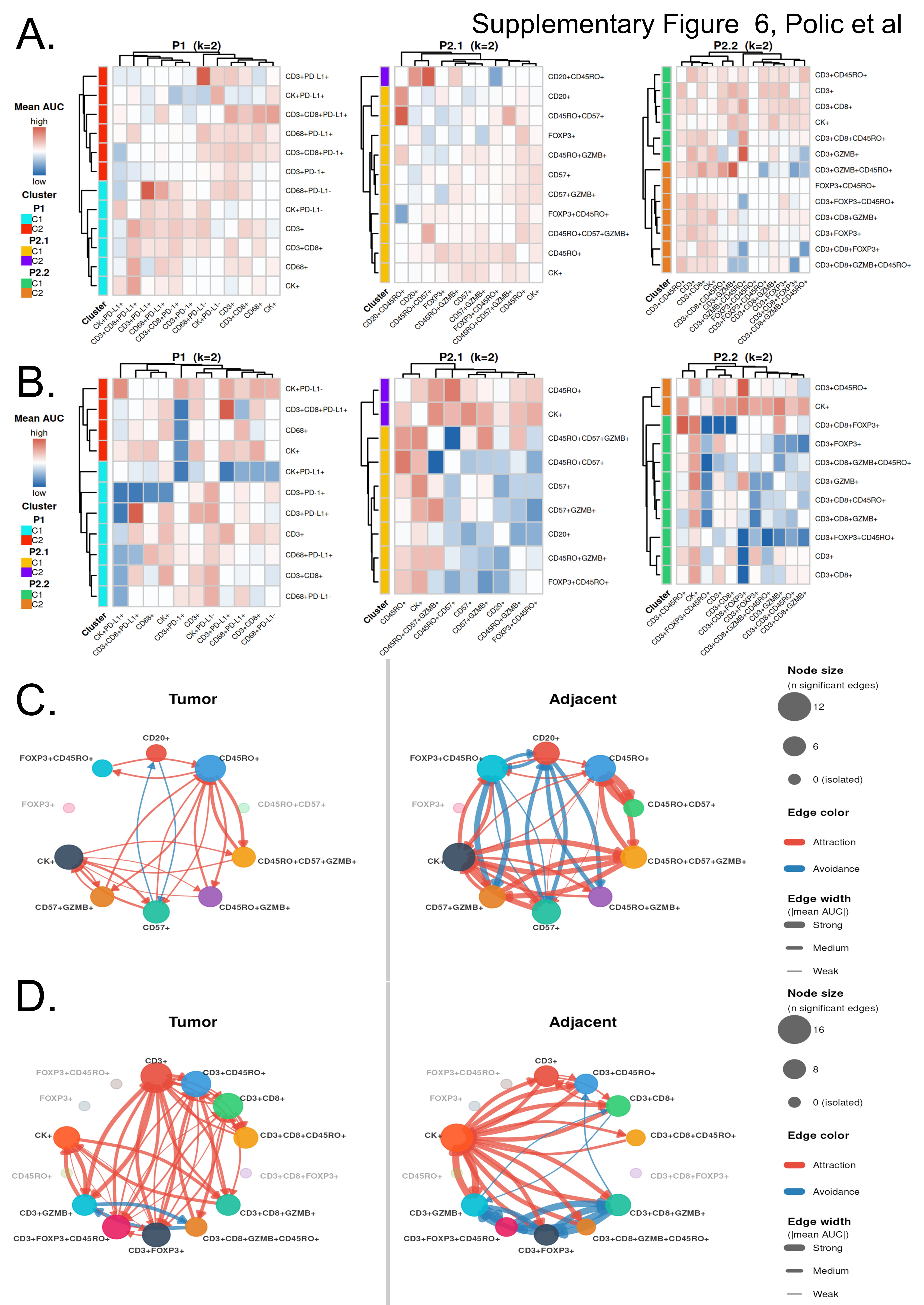
